# PacBio SMRT-based full-length transcriptome data of *Ascosphaera apis* mycelium and spore

**DOI:** 10.1101/2020.03.19.998625

**Authors:** Yu Du, Jie Wang, Huazhi Chen, Xiaoxue Fan, Zhiwei Zhu, Yuanchan Fan, Haibin Jiang, Cuiling Xiong, Yanzhen Zheng, Dafu Chen, Rui Guo

## Abstract

*Ascosphaera apis* is an entomopathogenic fungus that exclusively infects honeybee larvae, resulting in chalkbrood disease, a widespread fungal disease damaging the beekeeping industry all over the world. In this article, purified mycelia (Aam) and spores (Aas) of *A. apis* pure culture under lab condition were sequenced using PacBio Sequel platform. In total, 13,302,489 and 9,911,345 subreads were yielded from Aam and Aas, respectively; 394,142 and 274,928 circular consensus sequence (CCS) reads were identified as being full-length non-chimeric (FLNC) reads, with a mean length of 2820 bp and 2602 bp, respectively. Furthermore, 174,095 and 103,845 corrected isoforms were identified, with a N50 length of 3543 bp and 3262 bp, respectively. The reported full-length transcriptome data of *A. apis* mycelium and spore will provide a valuable resource for improvement of genome and transcriptome annotations as well as better understanding of transcript structure such as alternative splicing and polyadenylation.

**Value of the data:**

- Current dataset offers a set of high-quality full-length transcripts of *A. apis*.
- The data can facilitate the improvement of *A. apis* genome and transcriptome annotations.
- This dataset benefits further exploration of alternative splicing and polyadenylation of *A. apis* mRNAs.

## 1. Data Description

The shared full-length transcriptome data were from purified mycelia (Aam) and spores (Aas) of *A. apis* pure culture under lab condition. In total, 13,302,489 and 9911,345 subreads were respectively yielded from Aam and Aas, with an average read length of 1802 bp and 1742 bp, and an N50 of 3077 bp and 2731 bp (**Table 1**). Meanwhile, 464,043 and 315,135 CCS with a mean length of 2970 bp and 2733 bp were gained (**Table 1**). As presented in **Table 2**, for Aam and Aas, 402,415 and 277,919 reads were identified as being full-length (containing a 5’ primer, 3’ primer and the poly-A tail), and 394,142 and 274,928 were identified as being full-length non-chimeric (FLNC) reads with low artificial concatemers. The mean length of the FLNC reads was 2820 bp and 2602 bp, respectively (**Table 2**). As shown in **Table 3**, 182,165 and 107,906 unpolished consensus isoforms with a mean length of 2701 bp and 2461 bp were obtained; 121,776 and 70,701 high-quality isoforms as well as 58,307 and 35,946 low-quality isoforms were obtained after polishing these unpolished consensus isoforms with the Quiver algorithm. Finally, 174,095 and 103,845 corrected isoforms were identified, with a mean read length of 2728 bp and 2502 bp, and an N50 length of 3543 bp and 3262 bp (**Table 4**).

**Table 1.** Number and length distribution of PacBio SMRT sequencing

| Sample | Subreads num | Mean subread length | N50 | CCS | Mean CCS length |
| --- | --- | --- | --- | --- | --- |
| Aam[1] | 13,302,489 | 1802 | 3077 | 464,043 | 2970 |
| Aas | 9,911,345 | 1742 | 2731 | 315,135 | 2733 |

**Table 2.**
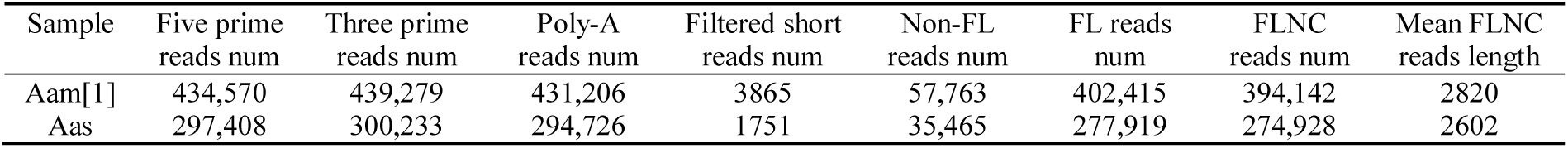
Statistics of PacBio SMRT sequencing output.

**Table 3.** Overview of FLNC reads clustering.

| Sample | Unpolished consensus isoforms num | Polished high-quality isoforms num | Polished low-quality isoforms num | Mean unpolished consensus isoform read length |
| --- | --- | --- | --- | --- |
| Aam | 182,165 | 121,776 | 58,307 | 2701 |
| Aas | 107,906 | 70,701 | 35,946 | 2461 |

**Table 4.** Summary of *A. apis* isoforms after correction.

| Sample | Total reads num | Length of total reads (bp) | Maximum length of total reads (bp) | Minimum length of total reads (bp) | Average length of total reads (bp) | N50 length of total reads (bp) | GC content of total reads (%) |
| --- | --- | --- | --- | --- | --- | --- | --- |
| Aam[1] | 174,095 | 474,928,820 | 13,808 | 50 | 2728 | 3543 | 49.00 |
| Aas | 103,845 | 259,853,381 | 14,756 | 57 | 2502 | 3262 | 49.04 |

## 2. Experimental Design, Materials, and Methods

### 2.1. Mycelia and spore sample preparation

*A. apis* was previously isolated from a fresh chalkbrood mummy of *Apis mellifera ligustica* larva [1] and preserved at the Honeybee Protection Laboratory of the College of Animal Sciences (College of Bee Science) in Fujian Agriculture and Forestry University. According to the previous method described by Jensen et al. [2] with some minor modifications [3], mycelium sampel and spore sample were respectively preparaed and frozen in liquid nitrogen, followed by storage at -80 °C until PacBio SMRT sequencing.

### 2.2. cDNA library construction and PacBio sequencing

The total RNA was extracted by grinding *A. apis* mycelia and spores in TRIzol reagent (Thermo Fisher, Shanghai, China) on dry ice according to the protocol provided by the manufacturer. The integrity of the RNA was determined using the Agilent 2100 Bioanalyzer (Agilent, USA) and agarose gel electrophoresis, and the purity and concentration of the RNA were detected using the Nanodrop micro-spectrophotometer (Thermo Fisher, Shanghai, China). Next, mRNA was enriched by Oligo (dT) magnetic beads and then reversely transcribed into cDNA using Clontech SMARTer PCR cDNA Synthesis Kit (TaKaRa, Shiga, Japan). The optimal amplification cycle number for the downstream large-scale PCR reactions was determined, and then used to generate double-stranded cDNA. Subsequently, >4 kb size selection was conducted based on the BluePippinTM Size-Selection System (Select science, Corston, UK) and equally mixed with the no-size-selection cDNA. Large-scale PCR was carried out for the next SMRT bell library construction; cDNAs were DNA damage repaired, end repaired, and ligated to sequencing adapters. Ultimately, the SMRT bell template was annealed to sequencing primer and then bound to polymerase, followed by sequencing on the PacBio Sequel platform using P6-C4 chemistry with 10 h movies by Gene Denovo Biotechnology Co. (Guangzhou, China). The raw data produced from PacBio SMRT sequencing were submitted to NCBI SRA database under accession number: SRR9887135 and SRR9887136.

### 2.3. SMRT reads processing and error correction

The raw sequencing reads generated form cDNA libraries of mecylium and spores were respectively classified and clustered into transcript consensus based on the SMRT Link v5.0.1 pipeline[4] supported by Pacific Biosciences. Briefly, (1) CCS (circular consensus sequence) reads were extracted out of subreads BAM file with minimum full pass of 1 and a minimum read score of 0.65; (2) CCS reads were classified into FLNC, non-full-length (nFL), chimeras, and short reads on basis of cDNA primers and poly-A tail signal, and reads shorter than 50 bp were discarded; (3) the FLNC reads were clustered by Iterative Clustering for Error Correction (ICE) software to generate the cluster consensus isoforms. The accuracy of PacBio reads was improved utilizing two strategies, firstly, the nFL reads were used to polish the above obtained cluster consensus isoforms using Quiver software to gain the FL polished high quality consensus sequences (accuracy≥99%); secondly, the low quality isoforms were further corrected using Illumina short reads produced from the same *A. apis* mecylium and spores samples using LoRDEC tool (version 0.8) [5].

### 2.4. Illumina short-read sequencing

Total RNA was respectively isolated from *A. apis* mycelia and spores using a Trizol Kit (Life technologies, USA). Then, poly-A mRNAs were isolated using Oligo (dTs) followed by fragmentation and reverse transcription with random primers (QIAGEN, Germany); second-strand cDNAs were synthesised using RNase H and DNA polymerase I. The double-strand cDNAs were purified using the QiaQuick PCR extraction kit (QIAGEN, Germany). Next, after agarose gel electrophoresis, the required fragments were purified using a DNA extraction kit (QIAGEN, Germany) and then enriched via PCR amplification in total volume of 50 μL containing 3 μL of NEBNext USER Enzyme (NEB, USA), 25 μL of NEBNext High-Fidelity PCR Master Mix (2×) (NEB, USA), 1 μL of Universal PCR Primer (25 mmol) (NEB, USA), and 1 μL of Index (X) Primer (25 mmol) (NEB, USA). The reaction conditions were as follows: 98 °C for 30 s, followed by 13 cycles of 98 °C for 10 s and 65 °C for 75 s, and 65 °C for 5 s. Finally, the amplified fragments were sequenced on the Illumina HiSeq 4000 platform (Illumina, USA) by Gene Denovo Biotechnology Co. (Guangzhou, China) according to the manufacturer’s protocols.

## Acknowledgments

This research was supported by the Earmarked Fund for China Agriculture Research System (No. CARS-44-KXJ7), the Science and Technology Planning Project of Fujian Province (No. 370 2018J05042), the Teaching and Scientific Research Fund of Education Department of Fujian Province (No. JAT170158), the Outstanding Scientific Research Manpower Fund of Fujian Agriculture and Forestry University (No. xjq201814), and the Scientific and Technical Innovation Fund of Fujian Agriculture and Forestry University (No. CXZX2017342, No. CXZX2017343).

